# Single-cell Analysis of Intracellular Transport and Expression of Cell Surface Proteins

**DOI:** 10.1101/2025.06.09.658704

**Authors:** Rekha Mudappathi, Vaishali Bharadwaj, Li Liu

## Abstract

Intracellular protein transport (ICT) is a tightly regulated process that orchestrates protein localization and expression, ensuring proper cellular function. Dysregulated ICT can lead to aberrant expression of surface proteins involved in cell-cell communication, adhesion, and immune responses, contributing to disease progression and therapeutic resistance. Cellular Indexing of Transcriptomes and Epitopes by Sequencing (CITE-seq) enables the simultaneous measurement of mRNA and surface protein levels in the same cell, providing a powerful opportunity to investigate the molecular mechanisms underlying surface protein regulation. In this study, we introduce a novel computational frame for Modeling Protein Expression and Transport (MPET) that evaluates the contribution of ICT activity to differential surface protein expression using CITE-seq data. MPET comprises three modules for identification of ICT– surface protein regulatory circuits across biological scales and their contributions to phenotypic variation. We applied MPET to analyze single-cell data from COVID-19 patients with varying disease severity. Our analysis revealed context-dependent recruitment of ICT genes and pervasive rewiring of ICT pathways throughout the course of disease progression. Notably, we found that even when the transcriptional levels of key immune response proteins remained stable, their expression on cell surface were significantly altered due to dysregulated ICT. MPET provides a valuable new tool for dissecting complex regulatory networks and offers mechanistic insight into post-transcriptional regulation of cell surface proteins in diseases.

## Introduction

Cell membranes exhibit a diverse array of proteins such as enzymes, transporters, ion channels, and receptors. These cell surface proteins actively contribute to a wide range of biological processes, such as signal transduction, cell adhesion, immune recognition, and molecular transport, playing vital roles in shaping cell functions ^1^. For instance, cluster of differentiation (CD) markers on the cell surface define cell types and cell differentiation stages ^2^ ^3^. Human leukocyte antigen (HLA), a key surface protein involved in immune responses, has been implicated in various health conditions when its expression is dysregulated ^4^. In cancer therapy targeting tumor microenvironment, several clinically approved drugs act on integrins, which are transmembrane adhesion receptors mediating cell-cell and cell-extracellular matrix interactions ^5^. Because of their distinct cellular locations and profound involvement in disease processes, cell surface proteins have emerged as promising diagnostic biomarkers and therapeutic targets ^6^ ^7^.

The expression of surface proteins is a complex process involving transcription, translation, post-translational modification, localization, and degradation ^8^. While numerous studies have focused on transcriptional regulation and shown that differentially transcribed mRNAs exhibit strong correlations with their protein products ^9^ ^10^, mRNA and protein abundance do not always align. Surface protein expression levels can be altered even when gene transcription levels remain unchanged ^11^ ^12^. This discrepancy underscores the need to investigate post-transcription regulatory mechanisms.

Notably, ICT is a critical process in ensuring proper protein transport, localization on cell membranes, and recycling, requiring precise coordination of multiple organelles and genes. As illustrated in **Fig. 1a**, it involves the translocation of proteins synthesized in ribosomes into the endoplasmic reticulum (ER), and subsequent transport through the Golgi apparatus for processing, modification, and sorting into secretory vesicles destined for the plasma membrane. In addition to direct transport to the plasma membrane, some proteins are routed through the endosomal-lysosomal network for recycling or degradation^13^ ^14^ ^15^ ^16^. Dysregulated ICT has been implicated in altered cell surface protein expression and associated with various diseases, including obesity, diabetes, cancer ^17^, and immune dysfunction in infections ^18^ ^19^. However, a comprehensive genome-wide understanding of ICT pathways involvement in surface protein regulation remains limited.

**Figure 1.**
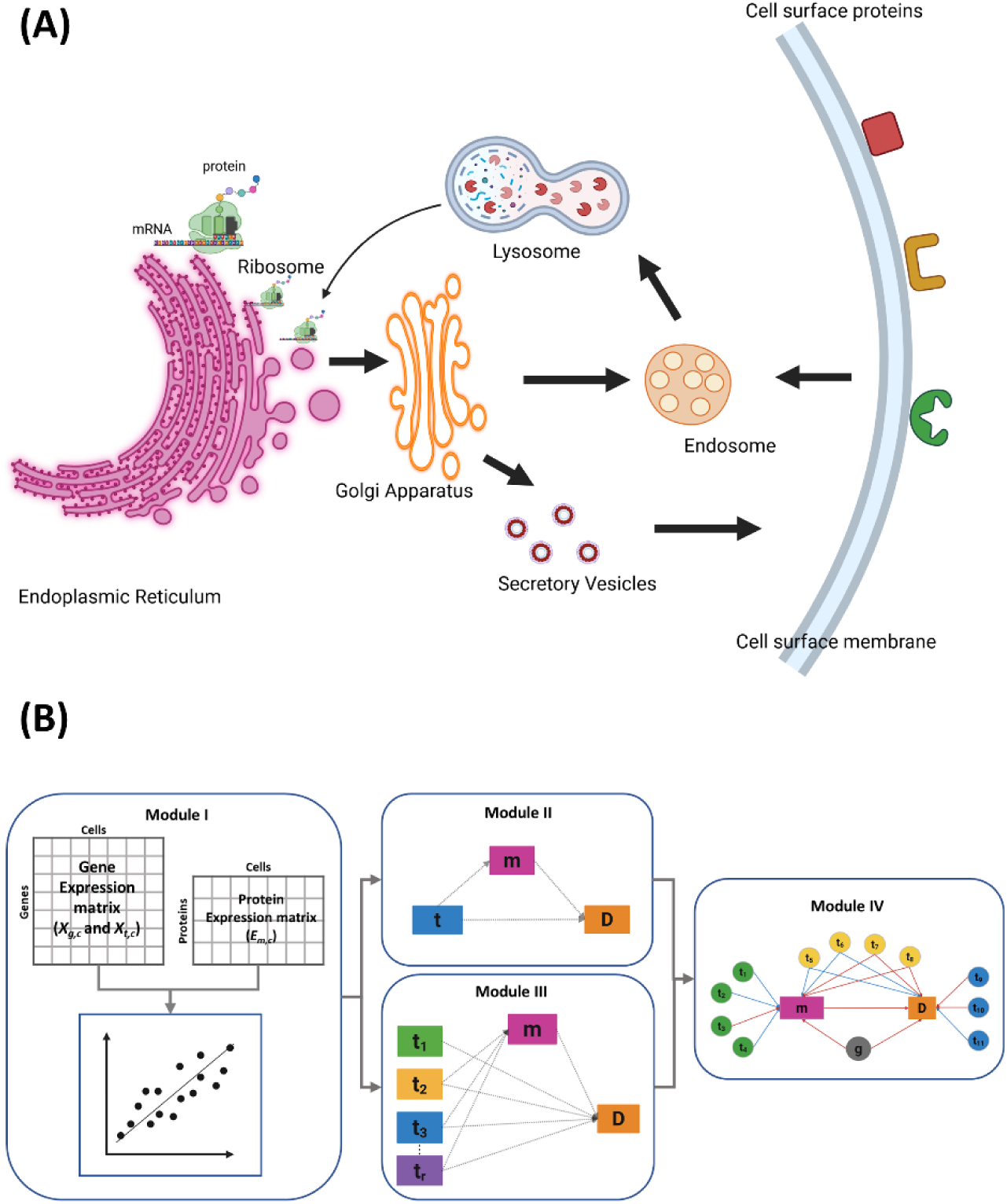
Overview of intracellular protein transport and MPET algorithm. **(A)** Ribosomes along with newly synthesized proteins attach onto endoplasmic reticulum (ER) membrane to enter ER lumen. After processing, membranous vesicles shuttle proteins from ER to Golgi apparatus. The proteins are then packed into secretory vesicles and transported to plasma membrane. Cells can internalize surface proteins via endocytosis. Endocytic vesicles fuse with recycling endosomes, from where they eventually move to lysosomes for degradation. Golgi cargo can also be sorted to endosomes, lysosomes, and ER for degradation and recycling. **(B)** CITE-seq data containing the gene expression matrix and protein expression matrix are input to MPET Module 1, in which mixed-effects regression analysis identifies ICT genes associated with surface protein abundance. The output from Module I can enter Module II for single-exposure mediation analysis or Module III for multiple-exposure mediation analysis. Module IV constructs a regulatory network to link surface proteins, their coding genes, and ICT genes to disease phenotype.

In this study, we present a novel computational approach for Modeling Protein Expression and Transport (MPET), to investigate regulatory circuits of surface protein expression in the context of ICT processes. This method leverages Cellular Indexing of Transcriptomes and Epitopes by Sequencing (CITE-seq) data, which simultaneously captures the single-cell transcriptome profile and expression level of selected surface proteins measured through Antibody-Derived Tags (ADTs) ^20^. Given a surface protein, MPET integrates the mRNA abundance of its initial transcripts, the ADT-based protein abundance on the cell membrane, and mRNA abundances of ICT-related genes. Using single-exposure and multi-exposure mediation models, it tests the hypothesis that ICT activity influences surface protein expression, which in turn affects disease phenotypes. Applying MPET to samples from COVID-19 patients, we observed pervasive disruption of ICT pathways, which led to aberrant expression of key immune response proteins even when the transcription of their coding genes remained unchanged.

## Results

### MPET Algorithm

We decompose the ICT regulatory network into small trios, each comprising of a surface protein *m*, its coding gene *g*, and an ICT gene *t*. MPET comprises four modules to examine the regulatory relationships within and between these trios.

Module I evaluates the association between the ICT gene and the surface protein within a trio. Given the CITE-seq data of a set of samples, we denote the expression level of a surface protein m in cell c from sample s as *E*_*m,c*_ measured by ADT abundance, and the transcription level of its coding gene g and an ICT gene t as *X_g,c_* and *X_t,c_*, respectively, measured by mRNA abundance. We test whether the ICT gene significantly regulates the expression of the surface protein after accounting for the transcription of its coding gene using a mixed-effects linear regression model,

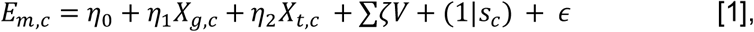

where η_0_ is the intercept, η_1_ and η_2_ are coefficients representing the fixed effect of *X*_*g,c*_ and *X*_*t,c*_, respectively, *V* is a set of covariates with corresponding ζ coefficients, 1|*s*_*c*_ represents the random effect accounting for multiple cells from the same sample, and ε is Gaussian-distributed errors. To correct for multiple comparisons, we adjust the nominal p values using the Benjamini-Hochberg method and calculate the false discovery rate (FDR). A trio with a significant non-zero η_2_ coefficient at FDR<0.05 is a putative transport trio (PTT).

Module II applies a single-exposure mediation model ^21^ to a PTT, testing whether the surface protein expression mediates the effect of an individual ICT gene on the phenotype, where the transcription level of an ICT gene *X_t,c_* serves as exposure, the abundance of a surface protein *E*_*m,c*_ acts as the mediator, and the phenotype *D*_*c*_ is the outcome (**Fig 2A**). These relationships are modeled as:

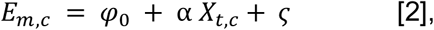

and

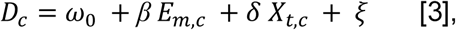

**Figure 2.**
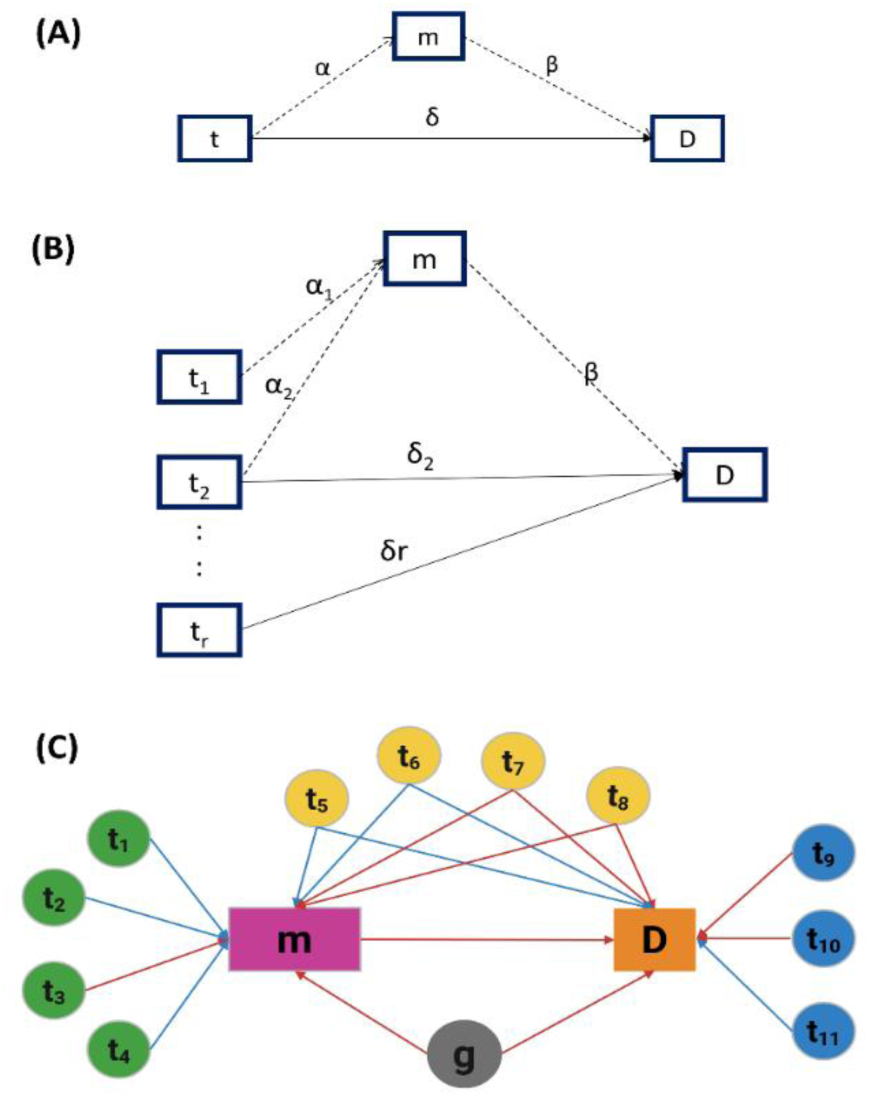
Mediation models in MPET. (**A**) Single-exposure mediation model. The transcription level of an ICT gene (t) may influence the disease phenotype (D) directly or indirectly via the expression level of a surface protein as a mediator (m). Coefficients in Eq. [2] and [3] are displayed along the corresponding edges. (**B**) Multiple exposure mediation model. Transcription levels of multiple ICT genes (t_1_,…,t_r_) are exposures. Coefficients in Eq. [4] and [5] are displayed along the corresponding edges. (**C**) Regulatory network represented as a directed acyclic graph. Nodes t_1_–t_4_ exhibit full mediation effects, Node t_5_–t_8_ show partial mediation. Nodes t_9_–t_11_ have null mediation. Red edges indicate positive associations, and blue edges indicate negative associations.

where φ_0_ and ω_0_ are intercepts, ⍺ and δ are coefficients representing the effects of ICT gene *t* on surface protein abundance and on phenotype, respectively, *ꞵ* represents the effect of surface protein abundance on phenotype, and ς and ζ are Gaussian-distributed errors. The total effect of ICT gene expression on phenotype is the sum of coefficients δ + ⍺ · *ꞵ*. Here, δ reflects the direct effect of the ICT gene on the phenotype, and ⍺ · *ꞵ* captures the indirect effect mediated through the surface protein. Significant non-zero indirect effects (⍺ · *ꞵ*) or direct effects (δ) with p-value < 0.05 were used to identify disease-associated ICT (daICT) genes.

We further classify the influence of daICT genes on phenotype into three categories: (i) full mediation (⍺ · *ꞵ* ≠0 and δ = 0), where the association between the ICT gene and phenotype is entirely explained through the surface protein mediator, (ii) partially mediation (⍺ · *ꞵ* ≠ 0 and δ ≠ 0), where both direct and mediated pathways contribute to the association, and (iii) no mediation (⍺ · *ꞵ* = 0 and δ ≠ 0), where the ICT gene affects the phenotype independent of surface protein expression.

Module III aggregates PTTs involving the same surface protein and uses a regularized multivariate mediation model ^21^ to evaluate the collective influence of multiple ICT genes on the surface protein abundance and the phenotype. Specifically, the abundance of a surface protein *E*_*m,c*_ in cell *c* acts as the mediator, the transcription levels of *r* ICT genes in the relevant PTTs, denoted as *X*_*t*=1…*r*,*c*_ serve as multiple exposures, and the phenotype *D*_*c*_ is treated as the outcome (**Fig 2B**). In the non-regularized form, the relationships are modeled as

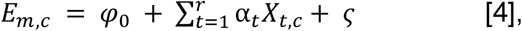

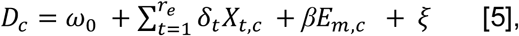

To prevent overfitting and reduce dimensionality, we apply *L1* regularization to both direct and indirect effects (**Methods and Materials**).

Module IV constructs a regulatory network (**Fig 2C**) as a directed graph, in which nodes represent a surface protein, its corresponding coding gene, daICT genes, and the phenotype, while edges depict regulatory relationships inferred from the mixed-effects model and mediation analysis (**Methods and Materials**).

Analysis of COIVD-19 data revealed concerted transportation of surface proteins.

We analyzed the CITE-Seq data set (GSE155673) from a previously published study of immunity in COVID-19 patients ^22^. This data set contains single-cell transcriptome profiles and abundances of 36 cell surface proteins in peripheral blood leukocytes from 12 age-matched individuals, including 5 healthy controls, 3 mild COVID-19 cases, and 4 severe COVID-19 cases. After quality control, normalization, and clustering, (**Methods and Materials**), we identified 13 distinct cell populations (**Fig 3A**), consistent with the original study, but with additional cell type annotations provided by our analysis. The original study reported significant transcriptional and surface protein changes in CD16+ monocytes from severe COVID-19 patients, characterized by increased activation, enhanced migratory capacity, and dysregulated antigen presentation ^23^. To investigate the role of ICT activities in these associations, we focused our analysis on CD16+ monocytes (5167 cells), which comprised 3,254 (10.7%), 632 (4.9%), and 1,281 (6.7%) cells in healthy control, mild case, and severe case samples, respectively.

**Figure 3.**
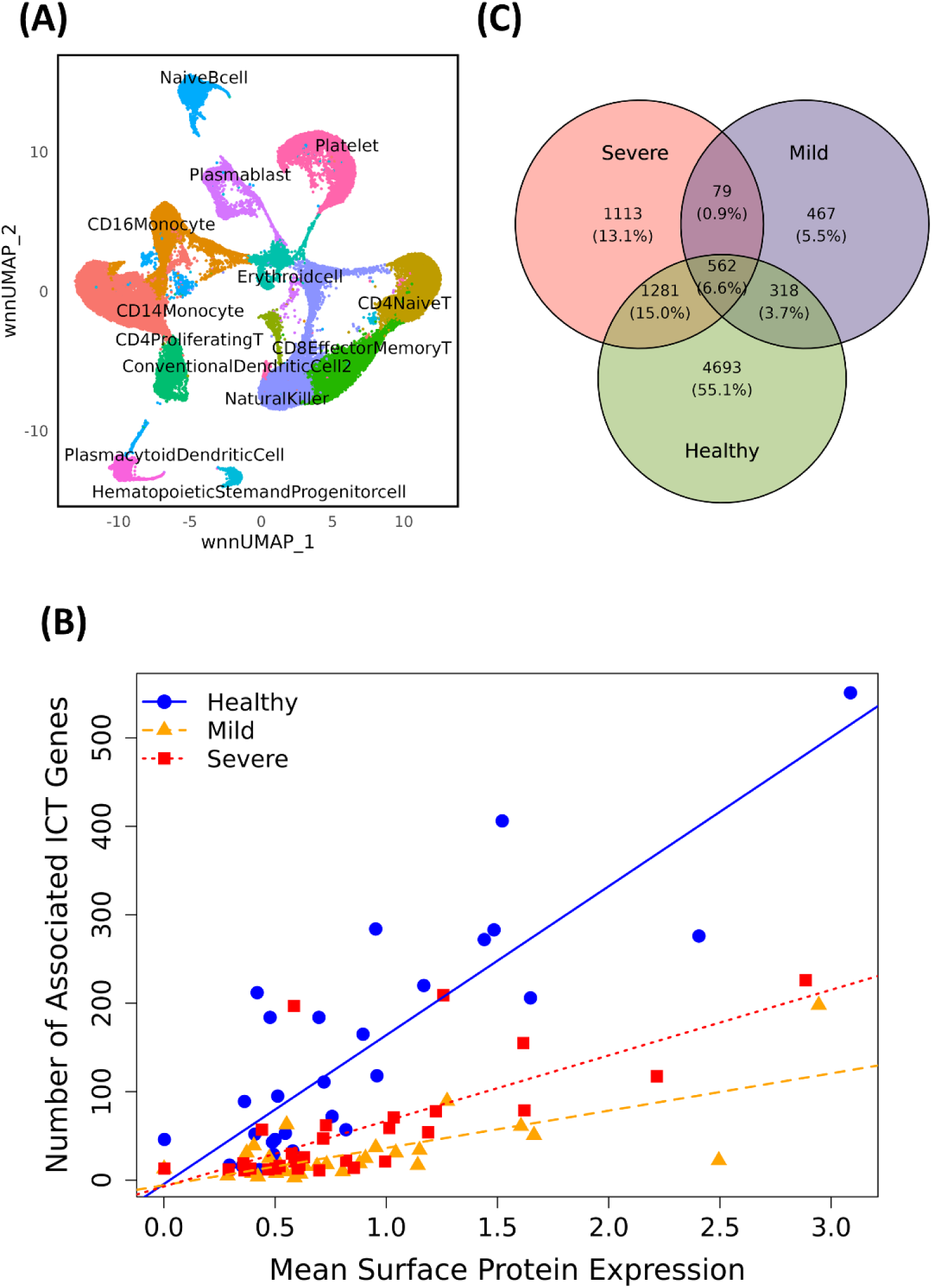
(**A**) UMAP visualization of CITE-seq cell clustering of cells from healthy control, and COVID-19 samples. (**B**) Scatter plot showing correlation between surface protein expression levels and the number of associated ICT genes in healthy cases, mild cases, and severe cases. (**C**) Venn diagram showing unique and shared PTTs between different phenotype groups.

### Concerted transport of surface proteins

Based on GeneOntology (GO) annotations, we identified 1839 ICT genes in the human genome (**Methods and Materials**). The CITE-seq dataset includes mRNA abundance data of 1,582 ICT genes and ADT abundance data of 30 surface proteins (**Table S4-5**). For surface proteins with multiple subunits, we included the coding gene for each subunit - specifically, *CD3D*, *CD3G*, and *CD3E* genes for the CD3 protein, *HLA-DRA, HLA-DRB1*, and *HLA-DRB5* genes for the HLA-DR protein, and *FCGR1A, FCGR2A, FCGR3A, FCGR1B, FCGR2B,* and *FCGR3B* genes for CD16 protein. In total, 39 surface protein─coding gene pairs, combined with 1,582 ICT genes, resulted in 61,695 unique trios for subsequent analysis.

Using MPET Module I, we performed mixed-effects regression analysis to identify PTTs. Because PTTs may vary across phenotypes, we performed this analysis for healthy controls, mild cases, and severe cases, separately. With a threshold of FDR<0.05, we identified 6,854, 1,426, and 3,035 PTTs in health controls, mild cases, and severe cases, respectively. All surface proteins and 80.5% (1,274) of ICT genes were included in at least one PTT. Notably, we found that highly expressed surface proteins recruited a greater number of ICT genes than those with low expression levels (Pearson correlation coefficient PCC=0.62, p-value = 7×10^-11^, **Fig. 3B**). This positive association was the strongest in cells from healthy individuals (PCC=0.84). Among all PTTs, only 562 (6.6%) were consistently identified across all three phenotypes, involving 175 unique ICT genes and 16 unique surface proteins (**Fig. 3C**). These observations suggest that the recruitment of ICT genes for transporting specific surface proteins is highly dynamic and context-dependent during disease progression.

Including ICT genes in the regression model also increased the explanatory power of coding gene transcription in surface protein expression. For each trio, we compared the full model used in Module I with a reduced model excluding the ICT gene. Across all surface protein–coding gene pairs, the full models yielded significantly higher η_1_ values (mean= 0.21 vs. 0.19, paired t-test *p* = 4×10^-3^), lower minimum p-values (mean= 0.06 vs. 0.14, *p* = 1×10^-2^), and greater mean R² values (mean= 0.05 vs. 0.04, *p* = 5×10^-14^) than the reduced model. These results suggest that incorporating ICT gene expression not only captures additional regulatory variation but also strengthens the association between coding gene transcription and surface protein abundance.

The η_2_ coefficient represents the effect of ICT gene transcription on surface protein expression. Based on η_2_ coefficients estimated in healthy individuals, we performed two-way hierarchical clustering, with rows representing surface protein-coding gene pairs and columns representing ICT genes. The resulting patterns confirmed the complex many-to-many relationships, showing multiple upregulating and downregulating ICT genes counterbalance each other to maintain the homeostasis of the expression of a surface protein. Meanwhile, an ICT gene could promote the expression of certain surface proteins while suppressing others (**Fig 4A)**. To examine if these clustering patterns based on η_2_ coefficients are due to correlation at the gene transcription level, we compared this dendrogram with that derived from pairwise correlations of coding gene mRNA abundance of the surface proteins **(Fig. 4B**). The resulting tanglegram revealed minimal concordance between the two trees supported by a Baker’s Gamma Index (BGI) of -0.03 and Cophenetic Correlation Coefficient (CCC) of -0.01, suggesting the hierarchical patterns captured by the two methods are uncorrelated. These results imply that transcriptional regulation and post-transcriptional regulation via ICT activities are likely two independent processes to fine-tune cell surface protein composition.

**Figure 4.**
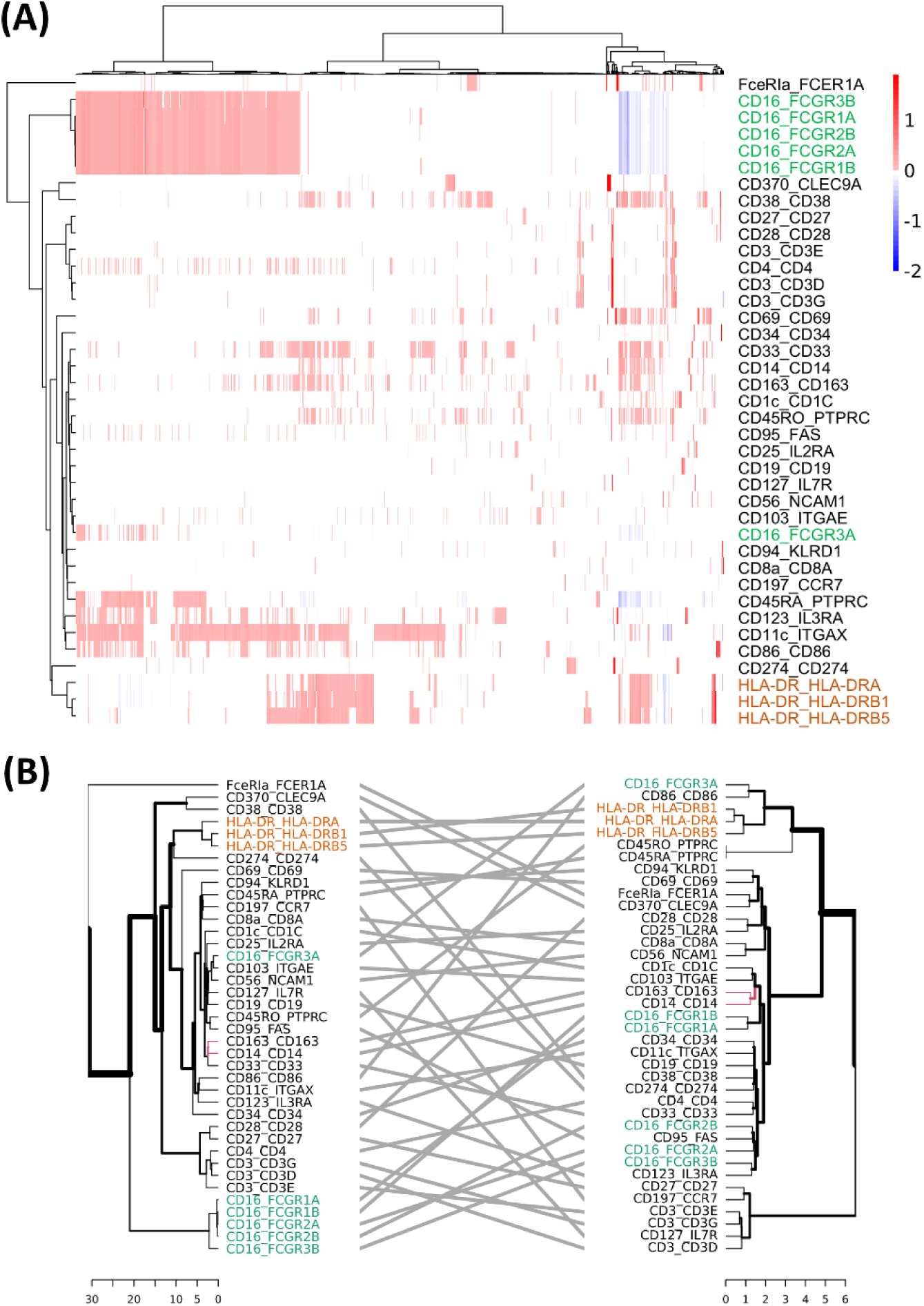
Surface protein and ICT association revealed in Module I analysis. **(A)** Heatmap showing association between surface protein-coding gene pairs and ICTs in Healthy group. **(B)** Tanglegram comparing hierarchical clustering structures based on ICT associations (left) and on coding gene transcription (right). HLA-DR subunits are labeled in orange and CD16 subunits in green in both panels.

We then examined surface proteins composed of multiple subunits and found a high degree of co-regulation. For example, HLA-DR consists of three subunits (labeled in orange in **Fig 4A** and **Fig. 4B)**. The effects of ICT genes on HLA-DR protein expression remained consistent regardless of which subunit’s coding gene was used in the Module I analysis. This concordance was also evident at the gene transcription level, suggesting that both transcriptional and ICT regulation of HLA-DR are synchronized at the protein complex level. In contrast, for the CD16 surface protein, which is composed of six subunits (labeled in green in **Fig 4A** and **Fig. 4B)**, only five coding genes are clustered together based on ICT effects. The sixth subunit, FCGR3A, showed distinct ICT associations and was positioned separately in the clustering results. Furthermore, the dendrogram based on gene transcription revealed greater dispersion among the six CD16 subunit genes, indicating more heterogeneous transcriptional regulation across subunits. These divergent patterns held true across severe and mild disease states as well (**Figure S1**).

### ICT-mediated mechanisms partially explain the discrepancy between gene transcription and protein expression of cell surface markers

Using the Wilcoxon rank-sum test implemented in the Seurat package, we identified 14, 19, and 5 differentially expression proteins (DEPs) between mild cases and health controls (MH), severe cases and health controls (SH), and severe and mild cases (SM), respectively, with FDR<0.05. Interestingly, for 4 (28.6%) of DEPs in MH, 7 (36.8%) in SH, and 1 (20.0%) in SM, their corresponding coding genes did not exhibit significant difference in mRNA levels across phenotype groups (all nominal p-value > 0.05, **Table S6)**, suggesting that dysregulation of these surface proteins may not be transcriptionally driven. For each of these DEPs, we applied the MPET Module II to investigate the contribution of ICT activity to the observed discrepancy.

One illustrative example is CD69, a well-known early activation marker on immune cells ^24^ ^25^. Prior studies have reported elevated CD69 expression in monocytes from COVID-19 patients, particularly in severe cases ^26^. Consistent with these findings, we observed that CD69 protein expression was significantly elevated in severe cases compared to healthy controls and mild cases (**Fig 5A**). However, the mRNA levels of *CD69* did not differ significantly between groups (nominal p-value =0.68) (**Fig 5B**). The MPET Module I analysis results showed that 145 ICT genes were involved in transporting CD69. We evaluated each of these ICT genes using Module II comparing severe cases against healthy controls and mild cases, which detected 138 (95.2%) daICT genes. In all models, CD69 protein expression was positively associated with disease status (β range: 0.16 - 0.49). An overwhelming majority (116, 84.1%) of daICT genes exhibited mediation effects and had positive α coefficients (0.02 - 1.07), indicating enhanced intracellular transport of CD69 (**Fig 5C**). These findings suggest that elevated ICT activity plays a substantial role in the overexpression of CD69 on cell surface, even in the absence of transcriptional changes of the CD69 coding genes.

**Figure 5.**
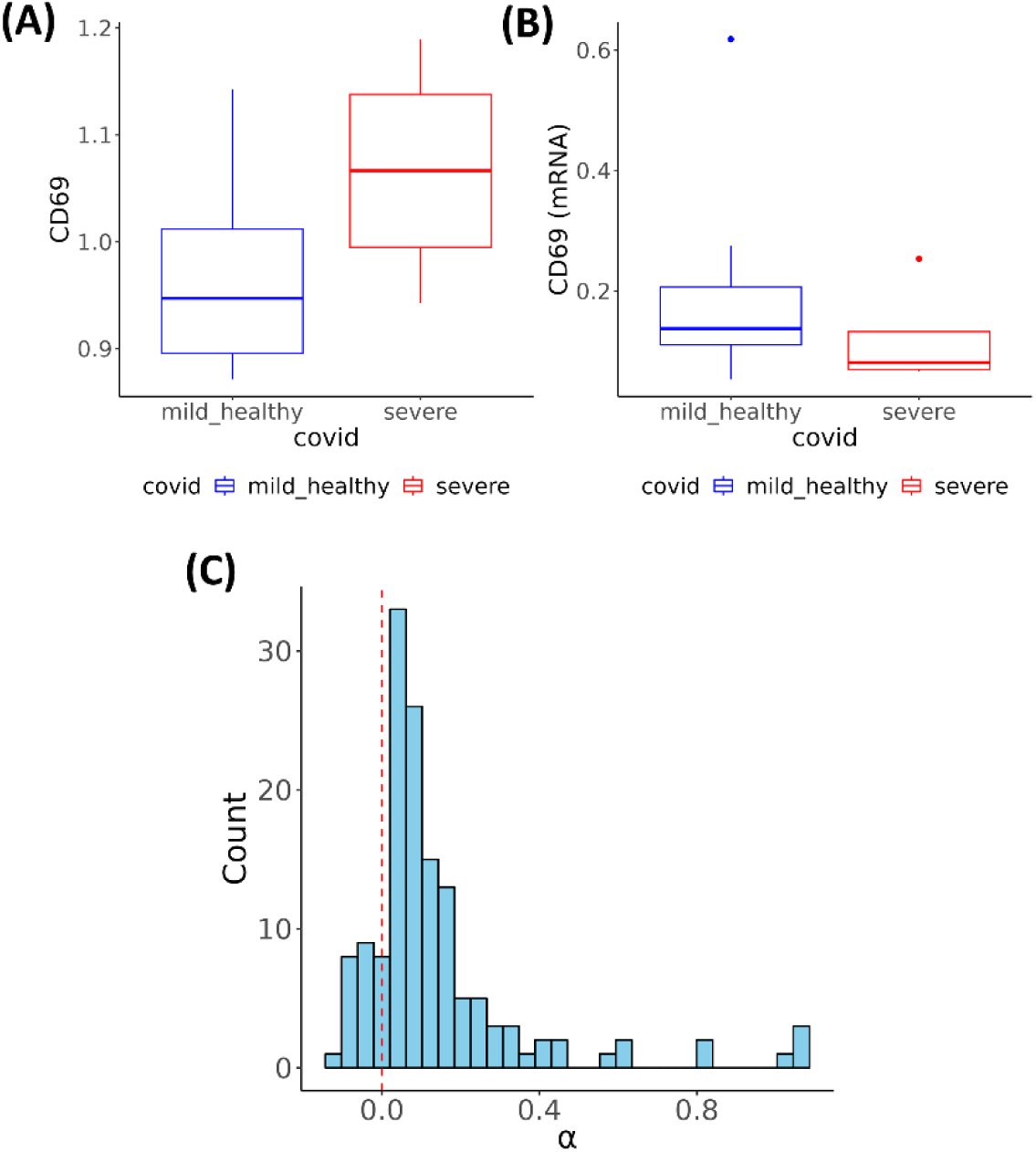
Modeling CD69 expression (**A**) Boxplot showing significantly increased surface protein expression of CD69 in CD16⁺ monocytes from patients with severe COVID-19 compared to mild and healthy controls (Wilcoxon rank-sum test, FDR < 0.05). (**B**) Boxplot showing no significant difference in *CD69* mRNA expression across the phenotype groups (nominal p-value = 0.68). (**C**) Distribution of the mediation effects (α coefficients) of daICT genes for the CD69 protein. One-sample t-test supports mean α >0 with p = 5.7 × 10⁻¹³.

### Multi-functional ICT genes contribute to aberrant expression of HLA-DR

HLR-DR is a key immune response protein, and low HLR-DR expression in monocytes is a strong predictor of poor COVID-19 prognosis ^27^ ^28^. Consistent with previous studies ^22^, we observed approximately 25% reduction in HLA-DR expression in severe COVID-19 cases compared to healthy controls and mild cases (p-value = 2×10^-19^ and 8×10^-21^, respectively, **Fig. 6A**). The Module I analysis identified 1,033 PTTs involving 526 unique ICT genes associated with HLA-DR. Among these, 382 showed strong effects (|η_2_| > 0.1) and were analyzed in Module III to evaluate their combined effects. At the optimal regularization parameter (λ=0.01), the multiple-exposure mediation model identified 152 daICT genes, including 61 (40.1%) with full mediation, 29 (19.1%) with partial mediation, and 62 (40.8%) with null mediation (**Fig 6B**). As expected, an overwhelming majority (53, 86.9%) of daICT genes with full mediation effects showed similar transcription levels between the disease groups. Furthermore, many (36 out of 61, 59.0%) were actively involved in transporting HLA-DR in healthy controls or mild cases, as captured by PTTs, but lost this function in severe cases, or vice versa. This observation suggested that disruption of ICT gene recruitment significantly contributes to disease progression, even when these ICT genes are normally transcribed.

**Figure 6.**
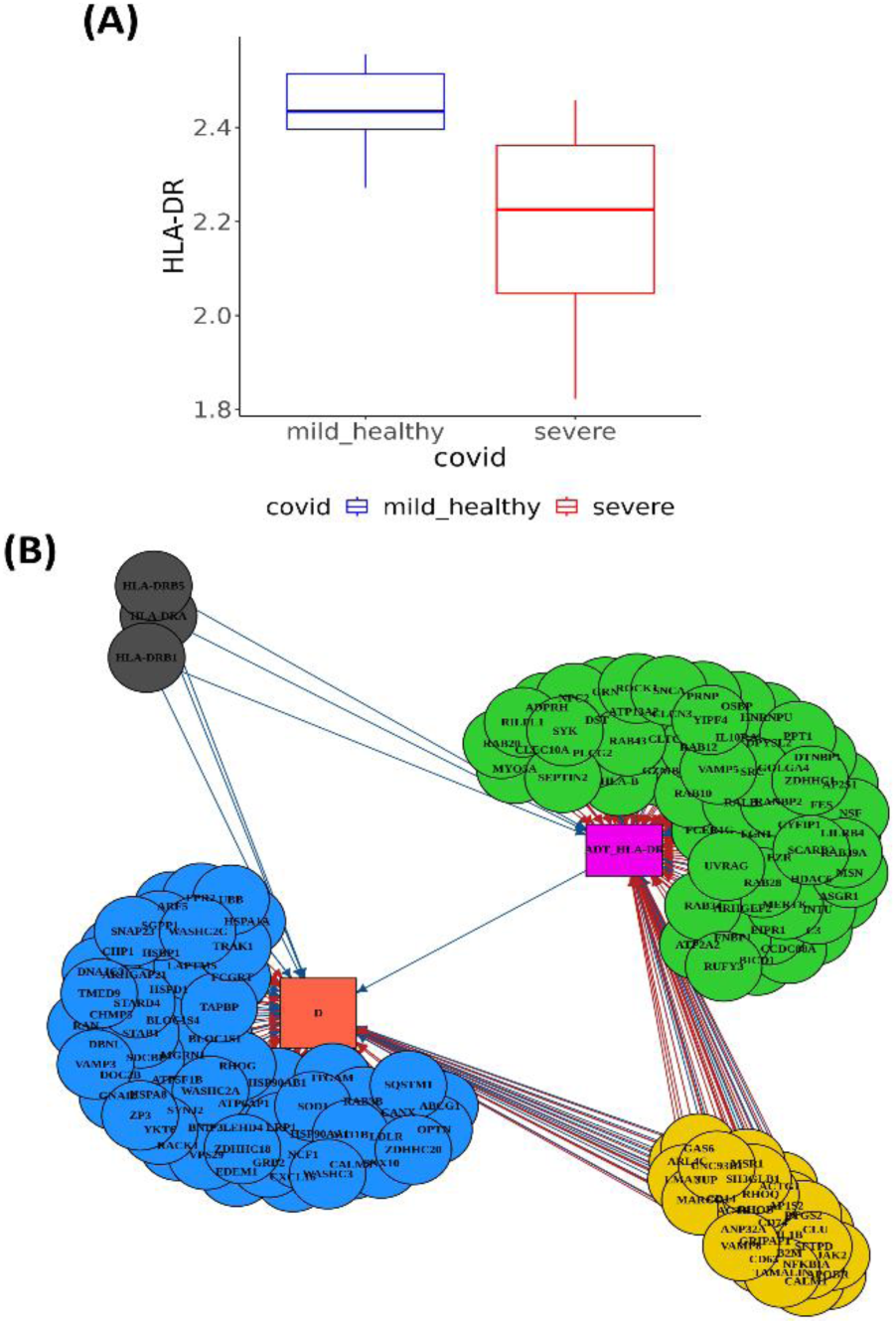
HLA-DR protein expression (**A**) Protein expression level is low in severe compared to healthy controls and mild. (**B**) Regulatory network with HLA-DR as the mediator, ICT genes as exposures, and disease severity (D) as the outcome. ICT genes are colored by mediation type - full mediation (green), partial mediation (yellow) or null mediation (blue).

We then annotated the 90 daICT genes with full or partial mediation effects based on GO annotations. Overrepresentation analysis of these genes against all ICT genes revealed a large number of significantly enriched GO categories (FDR<0.1, **Table S7**). Notably, more than half of these genes (46 out of 90) were related to endocytosis, enriched in GO categories such as positive regulation of endocytosis, receptor mediated endocytosis, endocytic vesicle, phagocytic vesicle, and lysosomal lumen, all of which are integral to the endocytic pathway. Among these endocytosis-related daICT genes, 16 showed partial mediation effects, influencing the phenotype both directly and through modulating surface protein expression. These genes include *CLU, CD14,* and *ACTB* that were significantly upregulated in severe cases (δ = 0.17, 0.03, and 0.08, respectively) and negatively associated with HLA-DR expression (⍺=-0.105, - 0.002, and -0.084, respectively) with FDR<0.05Meanwhile, *CD74, MARCO, MSR1*, and *VAMP8* were significantly downregulated in severe cases (δ range from -0.111 to -0.006) and positively associated with HLA-DR expression (⍺ range from 0.037 to 0.275). These patterns collectively suggest that endocytosis is overactivated in severe COVID-19, leading to reduced expression of HLA-DR on cell surface. Previous reports have shown that HLA-DR molecules can undergo recycling via endocytosis and become retained intracellularly in monocytes from patients with septic shock, contributing to reduced surface expression ^29^. Our findings suggest a similar mechanism in severe COVID-19 and further highlight the role of ICT genes in this process.

Several immune-related pathways were also enriched with daICT genes, including 39 (43.3%) participating in immune response, antigen presentation, cytokine signaling, and inflammatory responses. This reflects the fact that many ICT genes have multiple functions (i.e., moonlighting), serving roles beyond intracellular protein transport. These daICT genes encompassed a mix of pro- and anti-inflammatory genes, many of which were differentially expressed in severe cases compared to mild cases or healthy individuals. For example, *ACTG1, B2M, CD14, CD74, JAK2,* and *NFKBIA* are involved in the production and regulation of pro-inflammatory cytokines, such as interleukin-2, interleukin-6, and interleukin-8 ^30^ ^31^ ^32^ ^33^ ^34^ ^35^ ^36^. In contrast, *LILRB4,* and *VAMP8* are associated with anti-inflammatory functions ^37^ ^38^. Furthermore, certain genes, such as *CLU, GAS6, and GRN* are known to have dual roles, switching between pro- and anti-inflammatory functions depending on the cellular context ^39^ ^40^ ^41^ ^42^ ^43^. Given that severe COVID-19 is characterized by an imbalance between pro- and anti-inflammatory responses, often involving hyperinflammatory states (e.g., cytokine storm) and compensatory immunosuppression, the observed regulation of these daICT genes suggests that ICT genes influence the immune system at multiple levels ^44^ ^45^ ^46^ ^47^ ^48^.

These results collectively suggest that concerted alterations in the expression of multiple ICT genes contribute to the reduced surface expression of HLA-DR in severe COVID-19 cases. Many of these genes also play critical roles in immune regulation, potentially synergizing with the immunological functions of HLA-DR to drive immune suppression observed in severe disease.

### Comparison of single-exposure model and regularized multiple-exposure model

MPET supports two mediation models. The single-exposure model in Module II identifies daICT genes of a given surface protein, assuming that each ICT gene acts independently. To assess the combined effects of multiple ICT genes, Module III applies a regularized multiple-exposure model, accounting for potential correlations between ICT genes and penalizing those with weak effects.

To compare the two models, we applied Module II and III to each DEP in the severe vs healthy comparison of COVID-19 dataset and summarized the results (**Table 1**). On average, each DEP was associated with 181 daICT genes in the Module-II analysis, compared to only 85 daICT genes in the Module III analysis (**Fig. S2**). ICT genes exhibiting partial mediation effects constituted the majority (57.5%) of daICT genes in Module II results and were significantly more likely to be retained in the Module III results than those with full mediation effects (53.2% vs. 29.4%, Fisher’s Exact test *P=*2.2×10^-16^). However, among the retained ones, a substantial proportion converted from partial mediation to either full (20.1%) or null (49.9%) mediation. In contrast, while many ICT genes classified as having full or no mediation effects in Module II were dropped in Module III, those retained ones generally remained in the same category (67.3% for full mediation and 73.9% for no mediation).

**Table 1.** Comparison of mediation classifications between Module II and Module III.

| Module II /<br>Module III | Full | Partial | Nil | NA |
| --- | --- | --- | --- | --- |
| Full | 224 | 42 | 67 | 799 |
| Partial | 203 | 303 | 504 | 890 |
| Nil | 11 | 24 | 99 | 95 |
| NA | 25 | 0 | 22 | 0 |

As expected, ICT genes exhibiting strong effects tended to have consistent results between Module II and Module III analyses. Among daICT genes with full or null mediation in the Module II analysis, those retained in Module III results had effect sizes that were 68% to 73% higher than that of the dropped ones (mean |⍺. *ꞵ*| =0.094 vs. 0.056, t-test *p* = 2.6×10^-7^; mean |δ|=0.399 vs. 0.230, *p =* 2×10^-5^). For daICT genes with partial mediation in Module II, the retained genes had 41% to 50% higher effect sizes than those that were dropped (mean |⍺. *ꞵ*| =0.062 vs. 0.044, t-test *p =* 2×10^-13^; mean |δ|=0.310 vs. 0.206, *p =* 2×10^-16^). Moreover, for daICT genes that converted from partial mediation in Module-II to full mediation in Module-III, the direct effects were significantly weaker than those that did not convert (mean |δ| = 0.24 vs. 0.39; *p* = 7×10⁻⁹). Given these findings and the fact that the regularized model in Module III does not produce formal p values, effect size magnitudes and consistency between Modules II and III may serve as useful criteria for ranking and prioritizing candidate ICT genes for downstream validation.

## Discussion

MPET is a novel computational framework that models how ICT activities contribute to surface protein dysregulation through post-transcriptional mechanisms. The framework consists of three interconnected modules, each designed to test a distinct yet complementary hypothesis. Module-I filters ICT genes based on their associations with surface protein expression within the same phenotype group. Module-II applies a single-exposure mediation model to identify ICT genes that influence surface protein expression, which in turn affects phenotype, under the assumption that each ICT gene acts independently. Module-III employs a regularized multi-exposure mediation model to assess the combined effects of multiple ICT genes and prioritize those with the most substantial impact. Together, these three modules enable systematic identification of ICT-protein regulatory circuits and elucidate their contributions to phenotypic variation and disease progression.

To demonstrate the functionalities of MPET, we applied it to analyze single-cell CITE-seq data of CD16⁺ monocytes from COVID-19 patients. Across all phenotype groups, the analysis revealed multiple layers of regulatory complexity, showing that ICT gene recruitment is highly dynamic, context-dependent, and reprogrammed during disease progression. We found that surface proteins with higher abundance were associated with a greater number of ICT genes, suggesting an active recruitment process. Importantly, by incorporating ICT gene expression, MPET offers a mechanistic explanation for the frequently observed disconnect between mRNA and protein levels of surface proteins -demonstrating that regulation of ICT activity can modulate the expression of a surface protein independently of transcriptional changes in its coding gene. By classifying daICT genes into full and partial mediation categories, MPET enables the discovery of ICT genes that influence disease through both transcriptional regulation and their impact on surface protein transport, as well as those that act primarily through post-transcriptional mechanisms. These findings underscore that even non-differentially expressed ICT genes can exert significant regulatory effects, as rewiring of intracellular transport networks may selectively recruit or exclude specific ICT genes depending on the context of disease. Together, these results highlight the power of MPET to dissect post-transcriptional regulation and reveal its critical role in shaping surface protein expression and cellular phenotypes.

Our analysis of COVID-19 data pioneered the discovery of the significant roles of ICT activity in this disease. Specifically, we investigated the reduction of HLA-DR, a well-established hallmark of immune dysfunction in severe COVID-19 cases. For the first time, our results pointed to accelerated endocytosis as a potential contributor to the downregulation of HLA-DR surface expression. Moreover, the findings highlight the multifaceted roles of ICT genes, many of which are involved not only in general intracellular transport, but also in immune signaling and regulation, underscoring their dual contributions to both cellular logistics and immune response. These discoveries have important implications for understanding COVID-19 disease mechanisms and guiding therapeutic development. For example, our analysis identified *CLU* as a key regulator of several surface proteins, including HLA-DR CD14, CD38, CD16, CD69, CD11c, and CD123. In all models, *CLU* exhibited partial mediation effects, influencing disease severity both indirectly through its role in intracellular transport and directly through other pathways. Previous studies have shown that *CLU* is upregulated in infections caused by multiple coronaviruses ^49^, and has been linked to coagulation ^50^ and MHC class II-associated immune pathways ^51^. These results collectively suggest that *CLU* may serve not only as a phenotypic marker but also as a potential therapeutic target in COVID-19, positioned at the intersection of multiple cross-talking pathways.

Beyond COVID-19, the MPET framework offers a broadly applicable strategy for dissecting surface protein regulation across a wide range of disease contexts, including cancer, autoimmune disorders, and neurodegenerative diseases. In cancer, intracellular transport regulates key processes such as surface receptor trafficking, metabolic adaptation, and cell invasion during tumor progression ^17^. Applying MPET can reveal how dysregulated transport networks contribute to uncontrolled proliferation, metastasis, and immune evasion, offering new insights into tumor biology and potential therapeutic targets. Disruptions in intracellular trafficking are increasingly recognized as central to the pathogenesis of neurodegenerative diseases such as Alzheimer’s and Parkinson’s, contributing to impaired protein sorting, lysosomal degradation, and ER-Golgi transport, ultimately leading to toxic protein accumulation and neuronal dysfunction ^52^. Applying MPET could help elucidate how dysregulated intracellular trafficking contributes to neuronal dysfunction and pathogenesis across these disorders. Thus, MPET could enable mapping of transport defects across diverse cell populations, paving the way for new therapeutic strategies targeting intracellular trafficking pathways.

There are several limitations in MPET that present opportunities for future enhancement. First, although Module III supports the simultaneous assessment of multiple ICT genes, the use of regularization can lead to the exclusion of ICT genes with modest but potentially meaningful effects, resulting in false negatives. Our comparison between Modules II and III can serve as a useful guide for identifying and recovering these candidates, which primarily include ICT genes with full or null mediation when their indirect or direct effects are weak, and those with partial mediation when the direct effect is small. Second, as the number of ICT exposures increases, model fitting in Module III becomes more complex and computationally intensive, and convergence issues may arise, especially in the presence of multicollinearity. Third, single-cell sequencing data exhibit a nested structure with sample-level variation, which is accounted for in Module I but not in Modules II and III due to computational constraints. Ignoring this hierarchical structure may lead to biased effect estimates and inflated statistical significance. Fourth, MPET does not currently support a multiple-exposure, multiple-mediator, multiple-outcome model, even though ICT networks and interactions among multiple surface proteins likely contribute concurrently to disease phenotypes. Recent advances in machine learning, including regularization-aware neural networks, dimension reduction techniques, and graph-based modeling, offer promising solutions to address these challenges, and enhance MPET’s capacity to model complex, multi-dimensional regulatory landscapes in health and disease.

## Conclusion

MPET is a novel computational framework that models intracellular transport networks to understand the complex interplay between gene transcription, post-transcriptional regulation of surface proteins, and phenotype. It provides new opportunities to identify regulatory circuits that shape cellular phenotypes. By uncovering mechanisms beyond transcription, MPET holds promise for advancing precision medicine.

## Methods and Materials

### Construction of Trios in Module I

To construct trios, each comprising a cell surface protein, its corresponding coding gene, and an ICT gene, we first queried the GO annotations for biological process (BP) terms related to intracellular and vesicle-mediated transport, molecular function (MF) terms containing keywords such as protein carrier activity and chaperone binding, and cellular components (CC) involved in endocytic and exocytic pathways including the Golgi apparatus, lysosome, plasma membrane, and microtubules. The complete list of GO terms used in this search, along with the search process in the hierarchical GO structure across BP, MF, and CC, is provided in **Supplementary Methods** and **Tables S1–S3**. We define ICT genes as those that participate in these biological processes or molecular functions and are located in these specific cellular components. Given a surface protein measured in a CITE-seq experiment, we identified its coding gene from the HUGO Gene Nomenclature Committee (HGNC) database. For proteins with multiple subunits, all corresponding coding genes were retrieved. Each unique combination of a surface protein, its corresponding coding gene, and an ICT gene forms a trio.

### Regularization in Module III

To prevent overfitting and reduce dimensionality, we apply L1 regularization to both direct and indirect effects. The penalty function is:

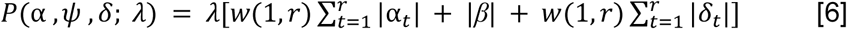

where λ is the regularization parameter, *w*(1, *r*) = *r*^1/4^ is the weight function to balance penalties for multiple ICT genes. We perform a grid search over a series of λ values ranging from 0.005 to 0.03, selecting the best model with the minimum Bayesian Information Criterion (BIC). As the coefficient *ꞵ* is not regularized, the impact of surface protein expression on phenotype is always retained in the model.

### Constructing Regulatory Networks in Module IV

To visualize the regulatory network involving surface proteins, ICT genes, and disease phenotypes, we constructed directed graph for a surface protein significantly associated with the phenotype. In each graph, the surface protein (m) and the phenotype (D) are always connected by an edge, reflecting their established association. Edges from daICT genes to the surface protein and disease status are drawn based on the nature of mediation. For daICT genes with full mediation, an edge is drawn from the ICT gene to the surface protein. For daICT genes with partial mediation, two edges are drawn, one from the ICT gene to the surface protein and the other from the ICT gene to the disease status. For daICT genes with null mediation, an edge is drawn from the ICT gene to the disease status. The coding gene of the surface protein is incorporated if its expression is significantly associated with surface protein abundance as assessed in Module I or if it is differentially expressed between disease groups. Edges are color-coded to reflect effect direction, with red for positive coefficients and blue for negative coefficients. This integrated regulatory network provides a comprehensive view of how a surface protein, its coding genes, and the associated daICT genes interact to influence disease-associated cellular processes.

### CITE-Seq preprocessing

The raw CITE-seq count matrices were preprocessed using Seurat (v4.1.2) ^53^. Cells with fewer than 100 detected genes, genes present in fewer than 5 cells, and cells with over 5% mitochondrial gene expression were filtered out. RNA expression was log normalized and z-transformed, while protein expression measured by ADT was centered log ratio normalized and z-transformed. Principal component analysis (PCA) was then applied, retaining 30 principal components for RNA and 15 for ADT. Clustering was done using Seurat’s weighted-nearest neighbor pipeline at a resolution of 0.2. Clusters with fewer than 5 cells were excluded. Marker genes for each cluster were identified, and cell types were annotated based on canonical marker genes ^54^.

### Differential Expression and Enrichment Analysis

Differential expression analysis for genes and surface proteins in the CITE-seq data was performed using the FindMarkers function implementing Wilcoxon rank-sum test in Seurat to identify differentially expressed genes (DEGs) and differentially expressed surface proteins (DEPs) between phenotype groups. Gene sets were tested for enrichment in GO-BP, GO-CC, and Hallmark pathways from the MSigDB database via overrepresentation analysis implemented in the R clusterProfile/enricher() function^55^.

### Comparison of Full and Reduced Models in Module I

For each surface protein–coding gene pair in healthy samples, we fitted a reduced model excluding the ICT gene and recorded the η₁ coefficient, p-value, and R² value. For the full model, we retained the maximum η₁ coefficient, the minimum p-value, and the mean R² value across all corresponding trios involving the same surface protein–coding gene pair. We then computed the average of these statistics across all pairs for both models. Compared to the reduced model, the full model yielded a significantly higher mean η₁ coefficient, lower mean p-value, and greater mean R² value. Paired t-tests confirmed that the differences were statistically significant, highlighting the added explanatory value of incorporating ICT gene expression.

## Availability of data and materials

The CITE-seq dataset analyzed in this study is publicly available from the Gene Expression Omnibus under accession number GSE155673^56^.

We also developed an R package implementing the MPET algorithm, which is freely available at: https://github.com/liliulab/MPET.

## Supplementary Methods

### GO Term Selection for ICT Gene Identification

To identify Gene Ontology (GO) terms associated with intracellular protein traffic, we used a targeted keyword strategy across the three GO domains: Biological Process (BP), Molecular Function (MF), and Cellular Component (CC). For BP, we selected GO terms containing keywords such as “intracellular transport,” “intracellular protein transport,” and “vesicle-mediated transport” to capture processes involved in intracellular trafficking and protein localization. For MF, we included terms matching “protein carrier activity,” “folding chaperone,” and “chaperone binding,” representing molecular functions linked to protein transport and stabilization. For CC, we identified terms associated with key trafficking organelles and compartments using keywords including “vacuole,” “autophago,” “endoplasmic,” “lysosome,” “Golgi,” “endosome,” “vesicle,” “vesicular,” “plasma membrane,” “intermediate compartment,” and “microtubule.” We then expanded each selected term to include all hierarchical descendants using the GO term ontology graph.

To ensure specificity for intracellular transport, we excluded GO terms unrelated to trafficking but potentially captured by overlapping keywords. These included terms related to phagocytosis, immune and neurosecretory granules, neurotransmitter exocytosis, and secretory processes such as histamine or protein secretion. Genes were retained only if they were annotated to both relevant GO terms in the BP or MF domains and localized to ICT-relevant compartments in the CC domain. This curated set of GO terms was used to define the ICT gene list for downstream analyses.

## Supplementary Figures

**Figure S1.**
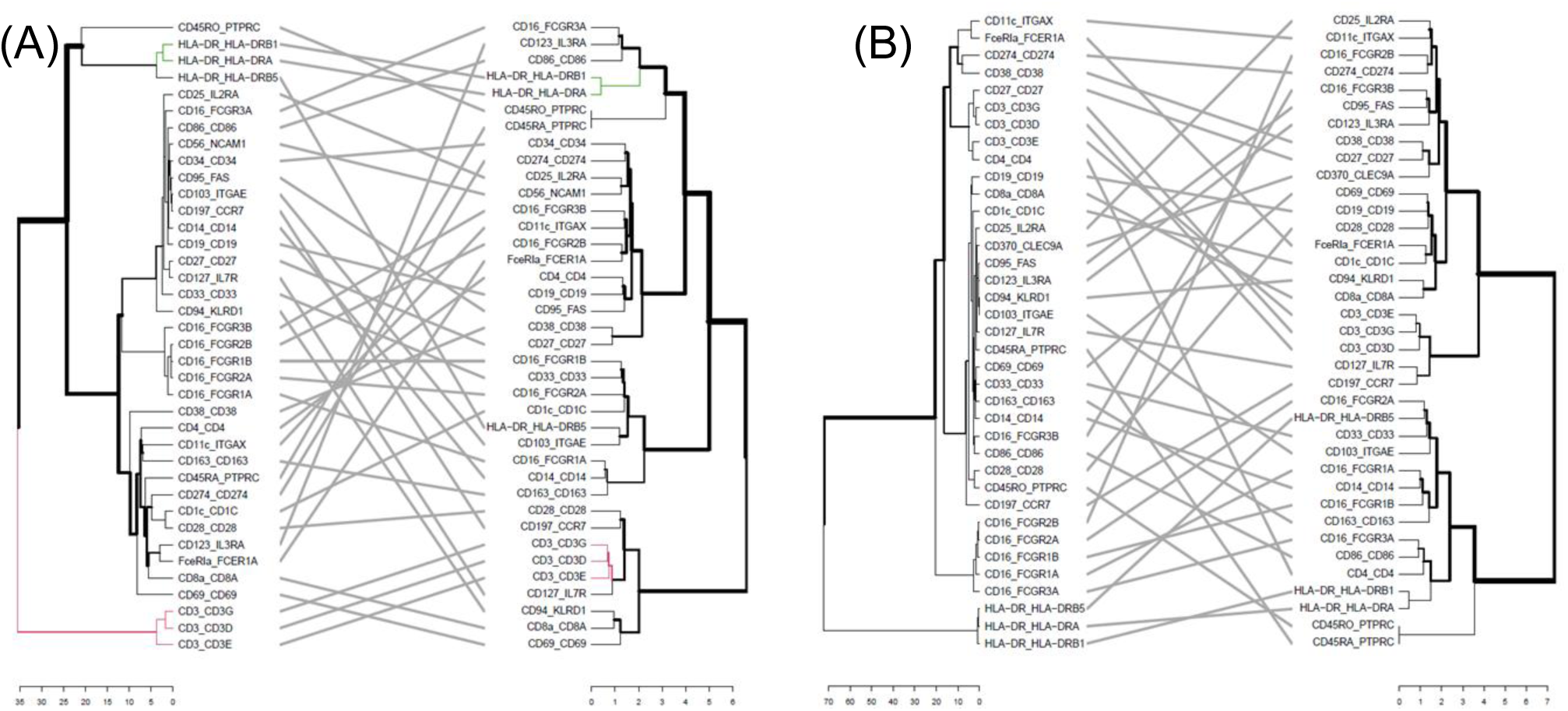
Comparison of dendrograms derived from η_2_ coefficients with those derived from pairwise correlations of mRNA expression for the coding genes in (A) Severe and (B) Mild COVID-19.

**Figure S2.**
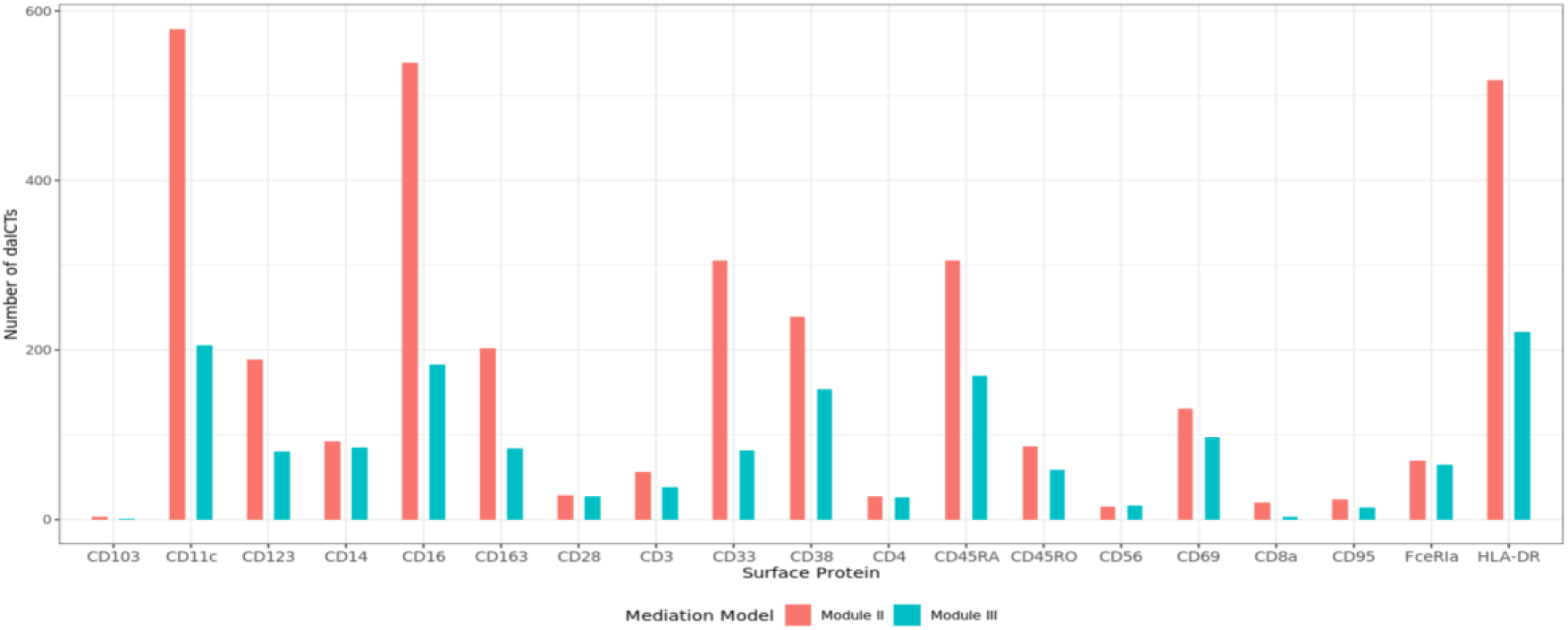
Comparison of the number of daICTs identified by Module II and Module III for each DEP in the severe vs. healthy comparison.

